# Whole-Brain Map of Top-down afferent Inputs to Nucleus Accumbens of the Mouse

**DOI:** 10.1101/2022.02.02.478577

**Authors:** Tonghui Xu, Jialiang Wu, Zhilong Chen, Zhao Li

## Abstract

Nucleus accumbens (NAc) is an important part of basal ganglia and receives major transsynaptic inputs from medial prefrontal cortex (mPFC), basolateral amygdala nucleus (BL) and hippocampus (HPC). The neural circuits formed by mPFC, BL, HPC with NAc are closely related to diverse brain functions such as decision-making, social behavior, reward seeking behavior and aggressive behavior. However, a lack of information about network structure of NAc top-down afference has limited our understanding of mechanisms of these functions. Here, we systematically analyzed whole brain patterns of inputs to mPFC/HPC/BL-NAc projectors via using (TRIO) labeling strategy. We found that the upstream input patterns between mPFC-core and mPFC-shell projections, BL-core and BL-shell projections, HPC-core and HPC-shell projections were similar. The projections from mPFC and BL to NAc receive a wide range of inputs, while the projections from HPC to NAc receive convergent inputs. Furthermore, all of mPFC/HPC/BL-NAc projectors receive considerable inputs from respective local regions. This study lays a foundation for further analysis of complex NAc functional mechanisms.

## Introduction

Nucleus accumbens (NAc) is a key component of basal ganglion^1^. It has been found that NAc integrates complex inputs to encode multiple signals and plays important roles in reward and punishment, feeding, spatial navigation, learning and memory via diverse projections^2-5^. Dysfunction of NAc is associated with many mental disorders, such as Alzheimer’s disease, drug addiction and depression^6-9^. Structurally, NAc consists of two subregions with very different functions: core and shell^10^. Specifically, the core plays an important role in instigating approach toward stimuli associated with rewards or safety. Whereas neural activity in the shell aids in inhibiting the emergence of behaviors, which may interfere with goal seeking^11^. The functional difference between NAc core and shell is due to respective complex anatomical connections^4^.

Our previous study showed that NAc mainly receives inputs from medial prefrontal cortex (mPFC), basolateral amygdala nucleus (BL) and hippocampus (HPC)^4^. As we know, these three brain regions play extensive and important roles in brain functions. The mPFC has been implicated as central to cognitive operation, attention, decision-making and working memory^12-17^. BLA is important in functions such as fear, anxiety, and reward^18-21^. HPC plays a pivotal role in memory and spatial navigation^22,23^. Additionally, in view of the diverse functions and complex projections of NAc, the complex circuits formed by mPFC, BLA, HPC with NAc has attracted wide attention. The mPFC-NAc circuit is essential for guiding individual behavior especially in situations requiring high mental concentration^24^. The BLA-NAc circuit is related to the individual’s response to reward and punishment^25^. Previous studies have shown that inhibiting the neural circuit of BL projection to NAc core or shell could weaken the individual’s response to reward or the punitive reward-seeking behavior, respectively^25^. The HPC-NAc circuit not only mediates foraging behavior^26^, but also helps animals use spatial information to guide discrimination methods and express preference for reward-related cues^27^. Nevertheless, the top-down afferent organization of these neural circuits remains unclear. We believe that revealing of the whole brain input patterns of mPFC/HPC/BL-NAc projectors is necessary to elucidate the circuits mechanism of their functions.

Here, we used the retrograde transsynaptic and rabies-based method TRIO to explore the whole brain input patterns of mPFC/HPC/BL-NAc projectors^28^.Our results reveal that mPFC/HPC/BL-NAc projectors receive a heterogeneous array of brain-wide afferents. All of the three projectors receive a high weight of local inputs.

## MATERIALS AND METHODS

### Animals

We used 8-10 weeks male C57BL/6J mice, C57BL/6J mice were purchased from Beijing Vital River (Beijing). All the experimental mice were reared in SPF mice room of Institute of life science, Nanchang University. All the mice were fed in standard cages with no more than 5 mice of the same sex in each cage. One male and two females were fed in one cage. The feeding conditions were as follows: ambient temperature 22 ± 2 °C, 12h circadian cycle, lighting period 7:00-19:00.

### Virus Information

All vectors and viruses used in this study were purchased from BrainVTA (BrainVTA Co., Ltd., Wuhan, China). The detailed production and concentration procedures for modified RV were previously described^29^. The final titer of RV-EnvA-ΔG-dsRed was 2 × 10^8^ infecting units per milliliter. For adeno-associated viruses (AAVs), AAV9-EF1a-FLEX-EGFP2a-TVA and AAV9-EF1a-FLEX-RG were packaged into 2/9 serotypes with final titers at 1–5 × 10^12^ genome copies per milliliter.

### Surgery and Viral Injections

We used 3% isoflurane to deeply anesthetize mice in oxygen, and 3D leveling. For tracing the experiments targeting mPFC/BL/HPC-NAc projectors, at day 1^st^, we mixed a small amount of 30nl AAV helper (AAV-EF1a-His-eGFP and rAAV2/retro-hsyn-Cre) at a ratio of 1:10 to inject NAc core and shell, respectively (core: A.P + 1.7 mm, M.L + 1.2 mm, D.V - 4.30 mm; shell: A.P + 1.7 mm, M.L + 0.45 mm, D.V −4.60 mm). At day 7^th^, we injected AAV9-EF1a-DIO-eGFP-TVA and AAV9-EF1a-DIO-RG (total volume: 50nl) at a ratio of 1:1 into mPFC, BL and HPC brain regions, respectively (mPFC: A.P + 1.94 mm, M.L + 0.50 mm, D.V - 3.00 mm; BL: A.P −1.70 mm, M.L + 3.00 mm, D.V −4.50 mm; HPC: A.P -3.40 mm, M.L + 3.20 mm, D.V - 4.00 mm). At day 21^th^, we injected 200nl RV-EnvA-ΔG-dsRed in same inject site into mPFC, BL and HPC brain regions. At day 28^th^, the mice brain was perfused. First, the mice were perfused with 0.01M phosphate buffered saline (PBS) buffer for 10 minutes in order to drain the whole blood; Then the mice were perfused with paraformaldehyde (PFA) in 0.01M PBS solution. After fixation, each sample was dehydrated with 10% sucrose solution, then we used melted agarose (Sigma-Aldrich, United States) to embedded each sample. The thickness of coronal brain slices was 50 μm by using a vibratome (Leica VT1000, Leica Microsystems). We used the method of taking one slice every other to collect the sample as the whole brain input data. The diluted glycerin solution was used to seal the film, and the film was stored in 4 °C refrigerator for later imaging.

### Imaging and analysis

The input neurons were imaged by VS120 slide scanner (Olympus, Japan) at 10× magnification. The starter cells were imaged by LSM 710 inverted confocal microscope (Zeiss) at 20× magnification. For each coronal sections, we adjust the exposure time and gain to ensure that the image quality is sufficient for cell counting. We used Image J to manually count all input neurons and starter cells according to a standard mouse brain atlas (PA). We used SPSS to analyze the data. The statistical tests included two tailed mismatch t test, Mann Whitney U test, one-way ANOVA analysis combined with Bonferroni correction and Kolmogorov Smirnov [K-S] test. All error lines are SEM.

## Results

### Strategies for tracing inputs to mPFC/BL/HPC-NAc projectors

To pathway-specific and rabies-based trace mPFC/BL/HPC-NAc projectors, we used TRIO method^28^ (**Figure 1A**): on the first day, we injected AAV-EF1a-His-eGFP and rAAV2/retro-hsyn-Cre into the core or shell of NAc of C57BL6/J mice (for representative examples, see **Supplementary Figure S1A**). One week later, we injected AAV helper (AAV9-EF1a-DIO-eBFP-TVA and AAV9-EF1a-DIO-RG) into mPFC, BL and HPC, respectively. The injected AAV helper only infected the neurons which projecting to NAc core or shell and expressing Cre recombinase. Cre recombinase was taken up by axonal terminals at the injection site and then was retrogradely transported to the somata. Two weeks after the injection of AAV helper, we injected RV-EnvA-ΔG-dsRed into mPFC, BL and HPC, respectively. RVG-deleted rabies virus (RV) could only infect the neurons with successful expression of TVA receptor. As we know, RV could not propagate retrogradely across synapses in the absence of RG. Therefore, only the mPFC, BL and HPC neurons which expressing TVA and RG can mediate the retrograde trans synaptic transmission of RV. These neurons simultaneously express blue fluorescence (AAV9-EF1a-DIO-eGFP-TVA) and red fluorescence (dsRed), and are identified as starter cells. The neurons which only expressing dsRed are the input neurons of mPFC/BL/HPC-NAc projectors. One week after the injection of RV, the mice were perfused.

**FIGURE 1.**
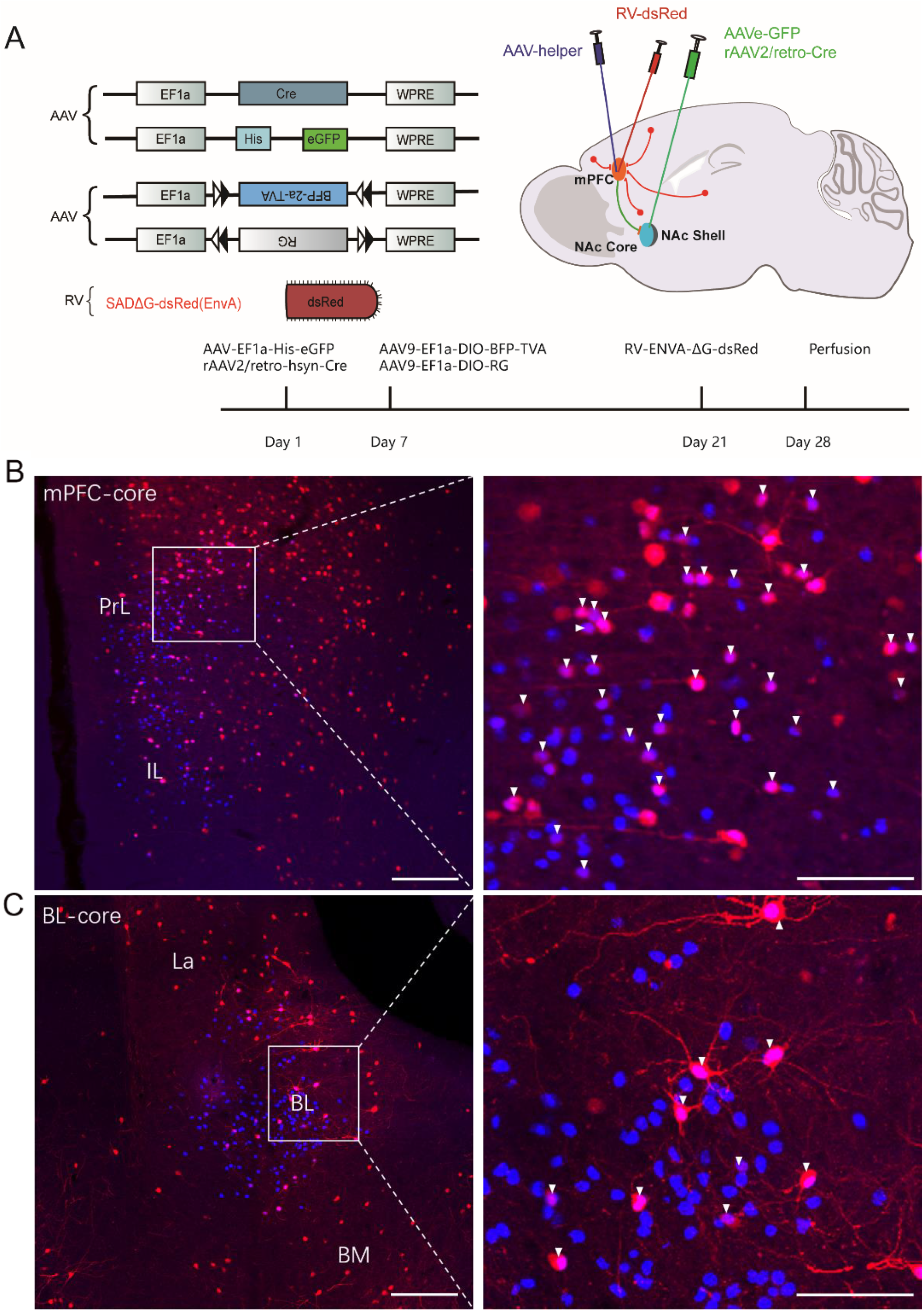
TRIO labeling strategy of mPFC/BL/HPC-NAc projectors. A. Virus carrier and injection flow chart. B. Left, representative coronal brain sections showing the location of starter cells in mPFC. Scale bar, 200 μm. Right, enlarged view of region in white box from the left image. Starter cells coexpressing BFP and dsRed, indicated by white arrowheads. Scale bar, 100 μm. C. Left, representative coronal brain sections showing the location ofx starter cells in BL. Scale bar, 200 μm. Right, enlarged view of region in white box from the left image. Starter cells coexpressing BFP and dsRed, indicated by white arrowheads. Scale bar, 100 μm. D. Left, representative coronal brain sections showing the location of starter cells in HPC. Scale bar, 200 μm. Right, enlarged view of region in white box from the left image. Starter cells coexpressing BFP and dsRed, indicated by white arrowheads. Scale bar, 100 μm.

To determine the accuracy of injection sites, we performed high-resolution imaging of the injection regions. We found that most of the starter cells in each group were restricted in the injection area, which indicated that the virus was accurately injected into the target sites and expressed successfully (**Figure 1B,C,D**). The results showed that both mPFC and HPC neurons which project to either NAc core or shell were dispersedly distributed in their respective regions (**Figure 1B,D**). Interestingly, most of the BL-core neurons were located in the ventrolateral part of BL (**Figure 1C**), while most of the BL-shell neurons were located in the dorsomedial part of BL. The reasonable speculation is that BL-core and BL-shell neurons belong to different neuron groups.

We counted and analyzed the relationship of input neurons and starter cells (n=24). The total number of input neurons we counted is 557084 (input neurons to mPFC-core: 33084, 38393, 30888, 27564; mPFC-shell: 42044, 59257, 20416, 24174; BL-core: 24487, 17496, 15612, 23644; BL-shell: 15374, 12385, 13644, 18745; HPC-core: 18523, 14085, 13562, 15395; HPC-shell: 18470, 17012, 22272, 20535) (**Figure 2A,B,C left and middle**). The number of starter cells ranged from 364 to 1894 (starter cells of mPFC-core: 1175, 1354, 877, 965; mPFC-shell:1544, 1894, 538, 647; BL-core: 705, 467, 413, 687; BL-shell: 441, 387, 364, 540; HPC-core: 671, 520, 364, 480; HPC-shell: 677, 589, 812, 764). Although injection volume of virus affects the number of labelled neurons, the numbers of transsynaptically labeled neurons had a linear relationship with the numbers of starter cells (**Figure 2A,B,C right**).

**FIGURE 2.**
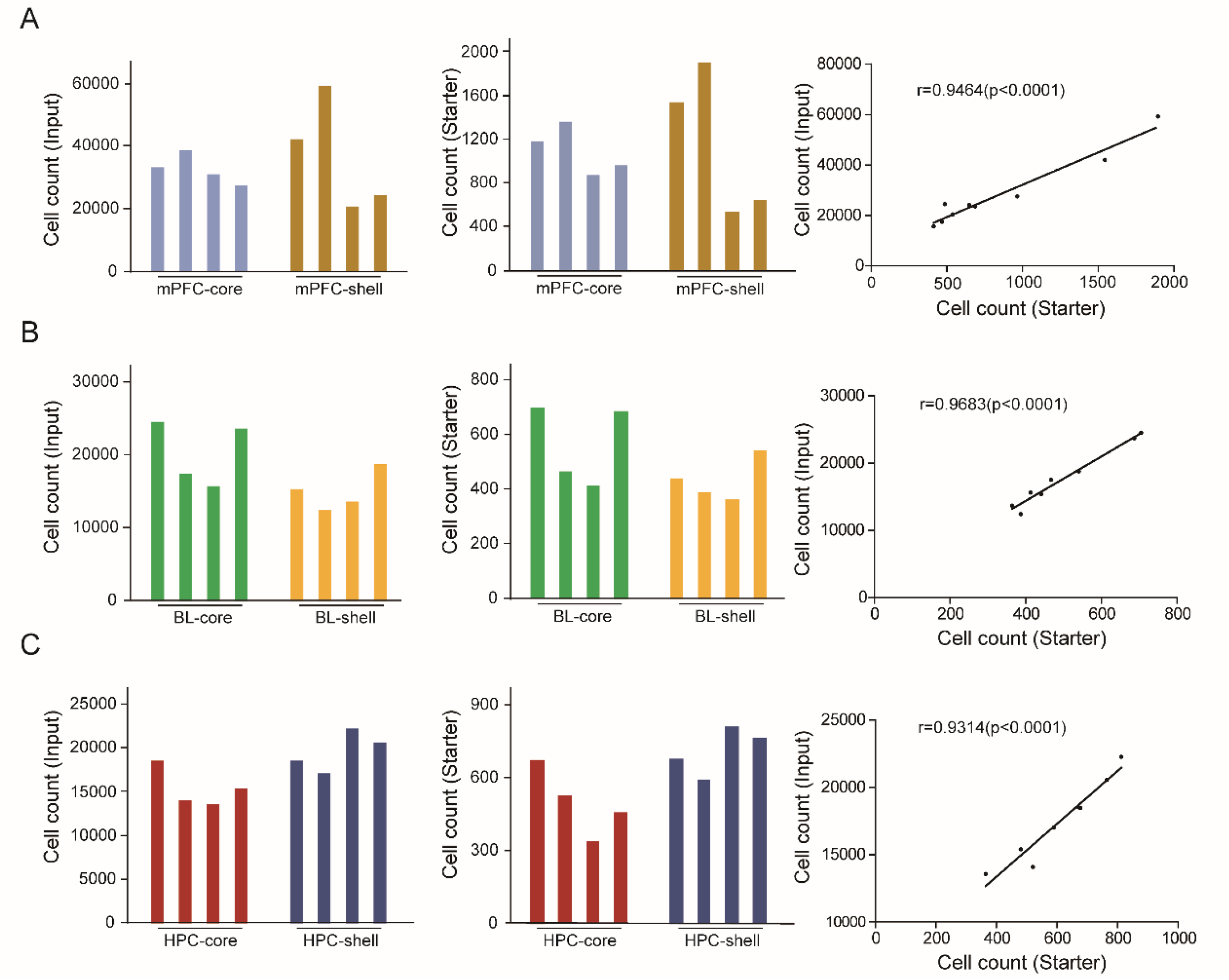
Statistical results of starter cells and input neurons of each sample. A. Left, the number of input neurons of mPFC-core and mPFC-shell. Middle, the number of starter cells. Right, a linear relationship was detected between the number of starter and input neurons. B. Left, the number of input neurons of BL-core and BL-shell. Middle, the number of starter cells. Right, a linear relationship was detected between the number of starter and input neurons. C. Left, the number of input neurons of HPC-core and HPC-shell. Middle, the number of starter cells. Right, a linear relationship was detected between the number of starter and input neurons.

### Whole-brain input patterns to mPFC/BL/HPC-NAc projectors

We systematically analyzed the brain-wide inputs of mPFC/BL/HPC-NAc projectors (**Figure 3**). Input neurons of mPFC/BL-NAc projectors were widely distributed throughout 114 brain areas. We studied distributions of their inputs in 64 cerebral regions and excluded the regions with input ratio less than 0.5% for the convenience of intensive study (**Figure 4**). Our results showed that inputs of HPC-NAc projectors are converged, while those of mPFC/BL-NAc projectors are highly dispersed (**Figure 4**). mPFC-NAc projectors received a high input weight from cortex, hippocampus and thalamus. The inputs from cortex to mPFC-NAc projectors were mainly concentrated in ventral orbital cortex (VO), medial orbital cortex (MO), infralimbic cortex (IL) of anterior cortex, prelimbic area (PrL), cingulate cortex (Cg) and secondary motor cortex (M2). mPFC-NAc projectors received local inputs (IL, MO, PrL, Cg) with a high input weight (**Figure 4, left**). We also found that there are 14 subregions throughout the thalamus project to mPFC-NAc projectors, especially the mediodorsal nucleus of the thalamus (MD) (**Figure 4, left**). BL-NAc projectors received projections from cortex, olfactory bulb, amygdala and hippocampus with a high input weight (**Figure 4, middle**). In cortex, BL-NAc projectors received inputs from posterior cortex with a high input weight, including the temporal association cortex (TeA), the auditory cortex (AU), the perinasal cortex (PRh), and the extranasal cortex (Ect). Notably, more than half inputs of HPC-NAc projectors came from CA1, subculum (S) and pyramidal cell layer of the hippocampus (52.96 ± 5.68%, **Figure 4, right**). Besides, in cortex, HPC-NAc projoctors received a high input weight from Ent (9.49%± 2.14%).

**FIGURE 3.**
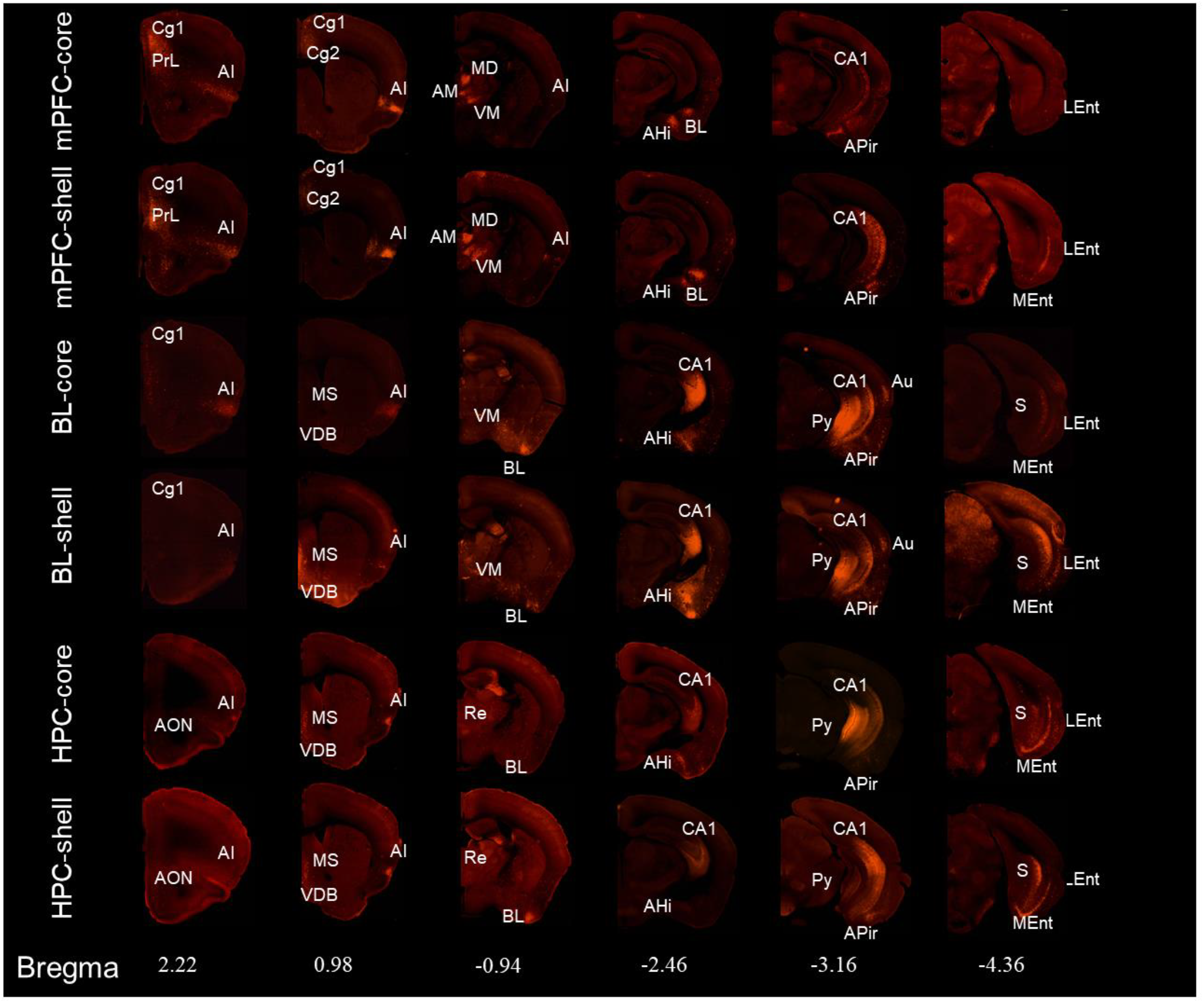
Representative coronal sections showed the upstream input distribution of mPFC/BL/HPC-NAc projectors (red, dsRed only, ipsilateral coronal section only). Scale bars, 1 mm.

**FIGURE 4.**
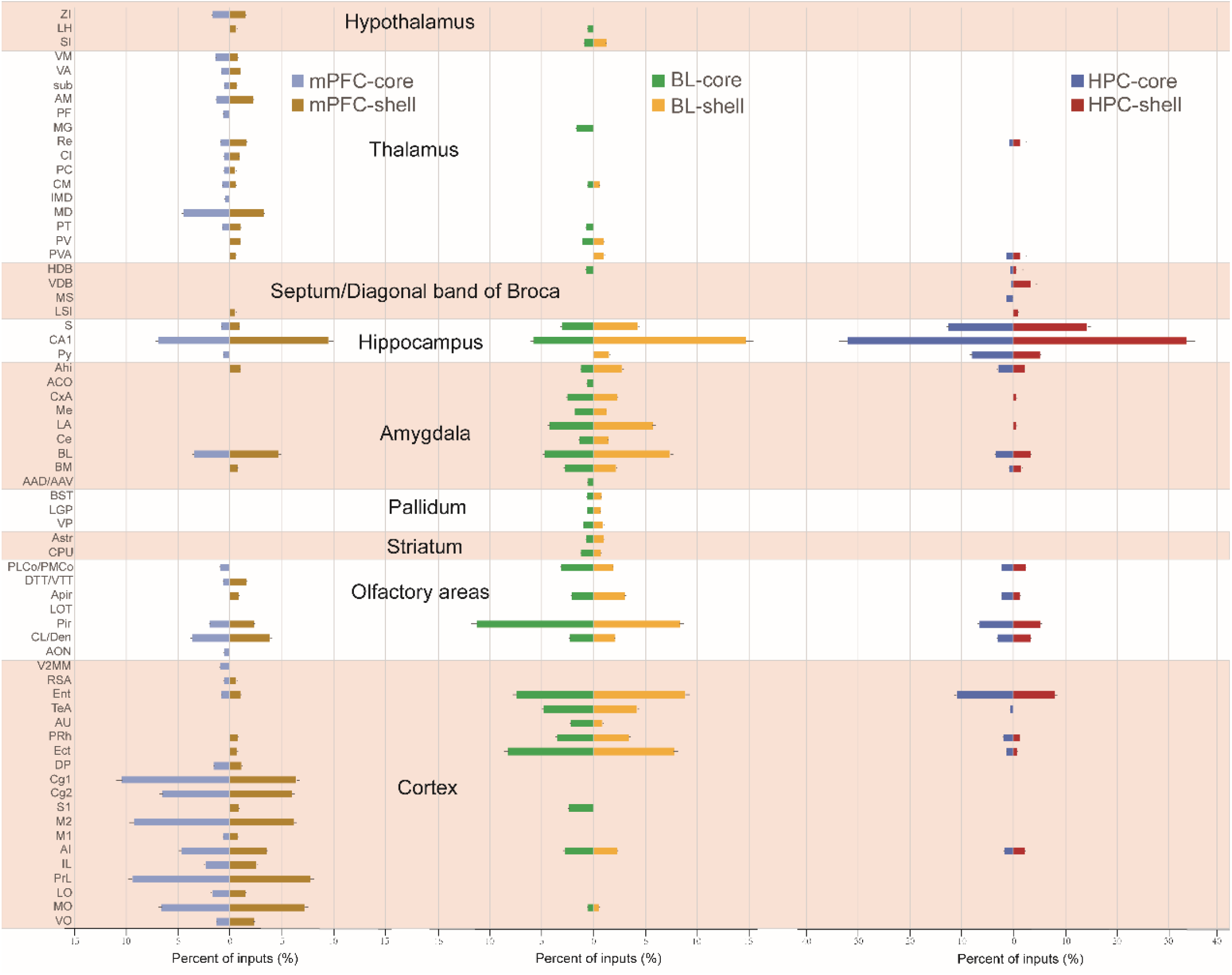
Quantitative analysis of the upstream input distribution of mPFC/BL/HPC-NAc projectors.

### Comparison between top-down afferences to NAc core and shell

The top-down afferent patterns between NAc core and shell were of little distinction. However, when the 64 cerebral regions we analyzed above were classed into 11 major brain areas, we found that there were significantly difference of top-down afference to several major areas between NAc core and shell. The cortex and amygdala provided more afferent inputs to mPFC-core than mPFC-shell, while the HPC and thalamic gave more inputs to BL-core than BL-shell (**Figure 5**).

**FIGURE 5.**
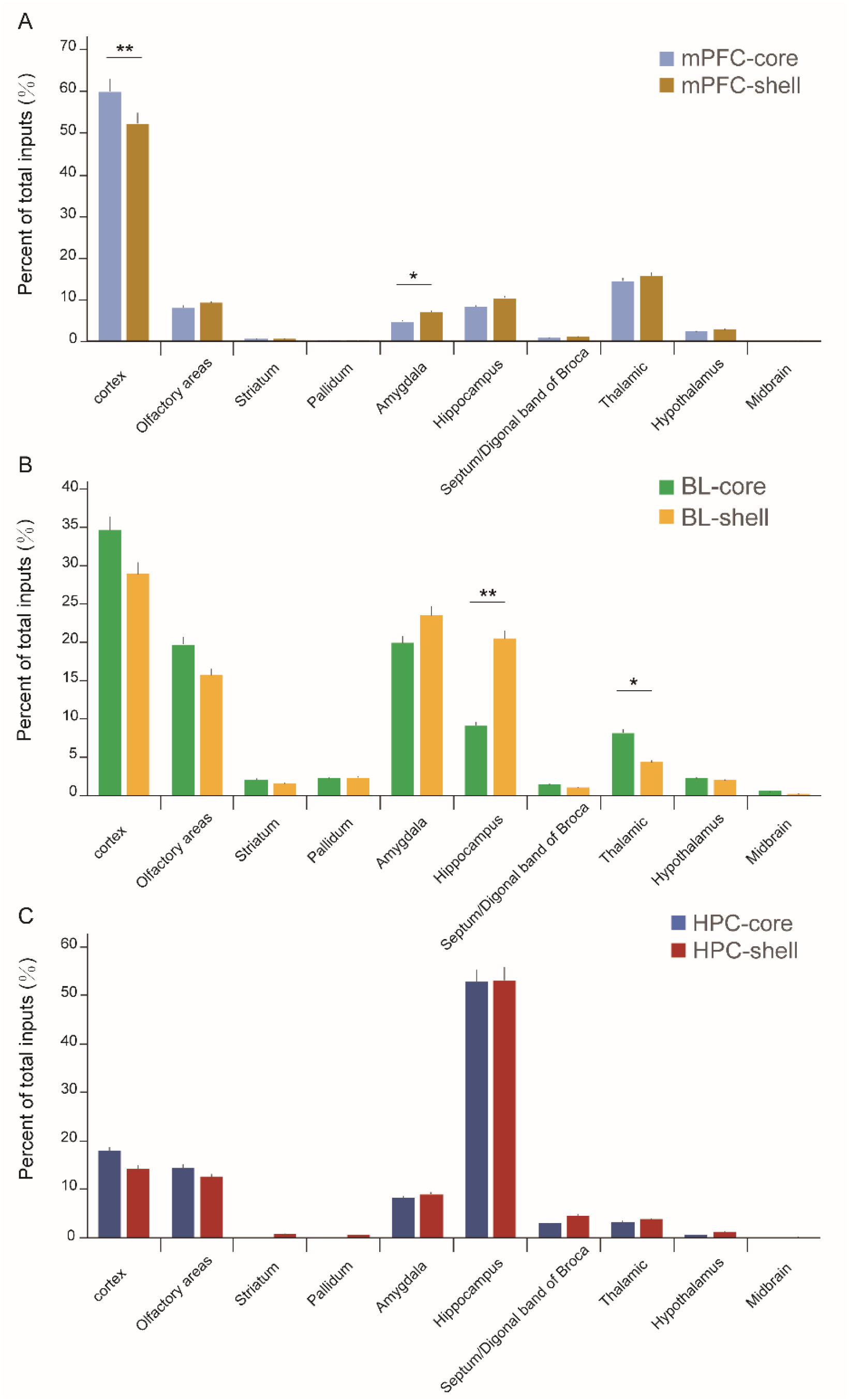
Distribution of input neurons of mPFC/BL/HPC-NAc projectors in the whole brain. A. Distribution of input neurons of mPFC-NAc projectors in the brain (2 experimental groups, 4 samples in each group). B. Distribution of input neurons of BL-NAc projectors in the brain (2 experimental groups, 4 samples in each group). C. Distribution of input neurons of HPC-NAc projectors in the brain (2 experimental groups, 4 samples in each group). Average value ± Standard error, * P < 0.01, * P < 0.05. One-way ANOVA combined with Bonferroni correction.

In order to quantify the similarities and differences in the top-down afferent patterns to NAc different subregions, we conducted a correlation analysis between inputs to mPFC-NAc core and mPFC-NAc shell, BL-NAc core and BL-NAc shell, HPC-NAc core and HPC-NAc shell. As showed in Figureure 6, each circle in the scatter plot represented one brain region (significant differences in red, *P* < 0.05), and the diagonal line represented the same input proportion for each pair. We found that the vast majority of the brain regions, represented by the open circles centered around the diagonals, provided similar top-down afferences to NAc core and shell, whether mPFC-core and mPFC-shell, BL-core and BL-shell, HPC-core and HPC-shell. In the mPFC-core and mPFC-shell targeted cases, only 7 of 104 brain regions provided significantly different inputs to mPFC-core and mPFC-shell (correlation coefficient, r = 0.939, *P* < 0.001; **Figure 6A**). As for the inputs to BL-core and BL-shell neurons, the correlation coefficient is 0.867 **(Figure 6B**), indicated BL-core and BL-shell neurons received more discrete inputs. The correlation coefficients between inputs to HPC-core and HPC-shell were extremely high (correlation coefficient, r = 0.986, *P* < 0.001; **Figure 6C**), probably due to large amounts of local afferences from HPC to HPC-NAc core and HPC-NAc shell neurons.

**FIGURE 6.**
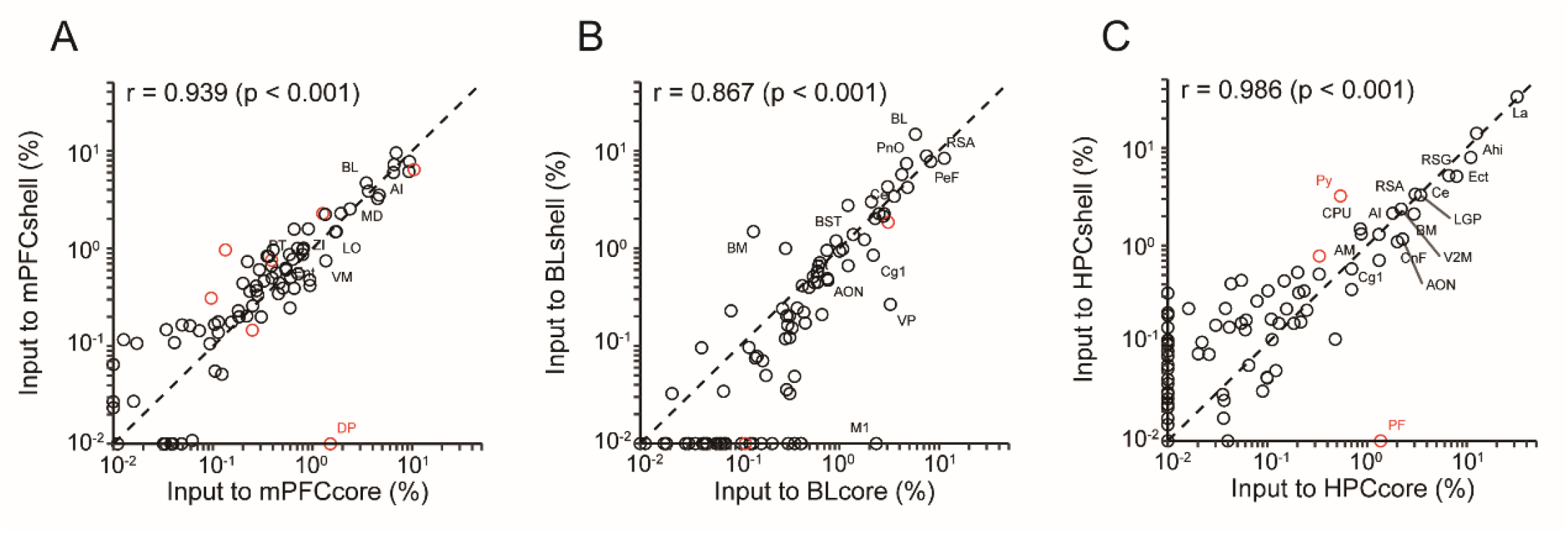
Comparison of input distribution among three groups. A. Comparison of input between mPFC-core and mPFC-shell neurons. B. Comparison of input between BL-core and BL-shell neurons. C. Comparison of input between HPC-core and HPC-shell neurons. Values are the means of the percent of total inputs from each region. Red circles indicate significant differences (p < 0.05, two-tailed unpaired t-test or Mann–Whitney U test). r, Pearson’s correlation coefficient.

### Comparison among the top-down afferences from Cortex, Amygdala and HPC to mPFC/BL/HPC-NAc projectors

As showed in Figureure 6, cortex, amygdala and HPC account for high proportion of top-down inputs to either NAc core or shell. Therefore, it is necessary to analyze the inputs from subregions of cortex, amygdala and HPC to mPFC/BL/HPC-NAc projectors in detail. Firstly, we found that cortex, amygdala and HPC account for high proportion of inputs to all of mPFC/BL/HPC-NAc projectors (coretex, 57.76%± 6.82%, 41.47%±4.82%, 21.87%±2.41%; amygdala, 12.53%±1.24%, 21.63%±2.76%, 8.51%±0.87%; HPC, 9.36%±0.94%, 14.70%±2.12%, 52.57%±4.33%, respectively). By comparison, cortex provided more inputs to mPFC-NAc projectors than to BL-NAc and HPC-NAc projectors. Interestingly, the top-down afferences from different subregions of cortex to NAc showed virtually preference. Specifically, mPFC and motor cortex only provided inputs to mPFC-NAc projectors, while extranasal cortex (Ect), perinasal cortex (PRh), auditory areas (AU), temporal association cortex (TeA) only provided inputs to BL-NAc and HPC-NAc projectors (**Figure 7B**).

**FIGURE 7.**
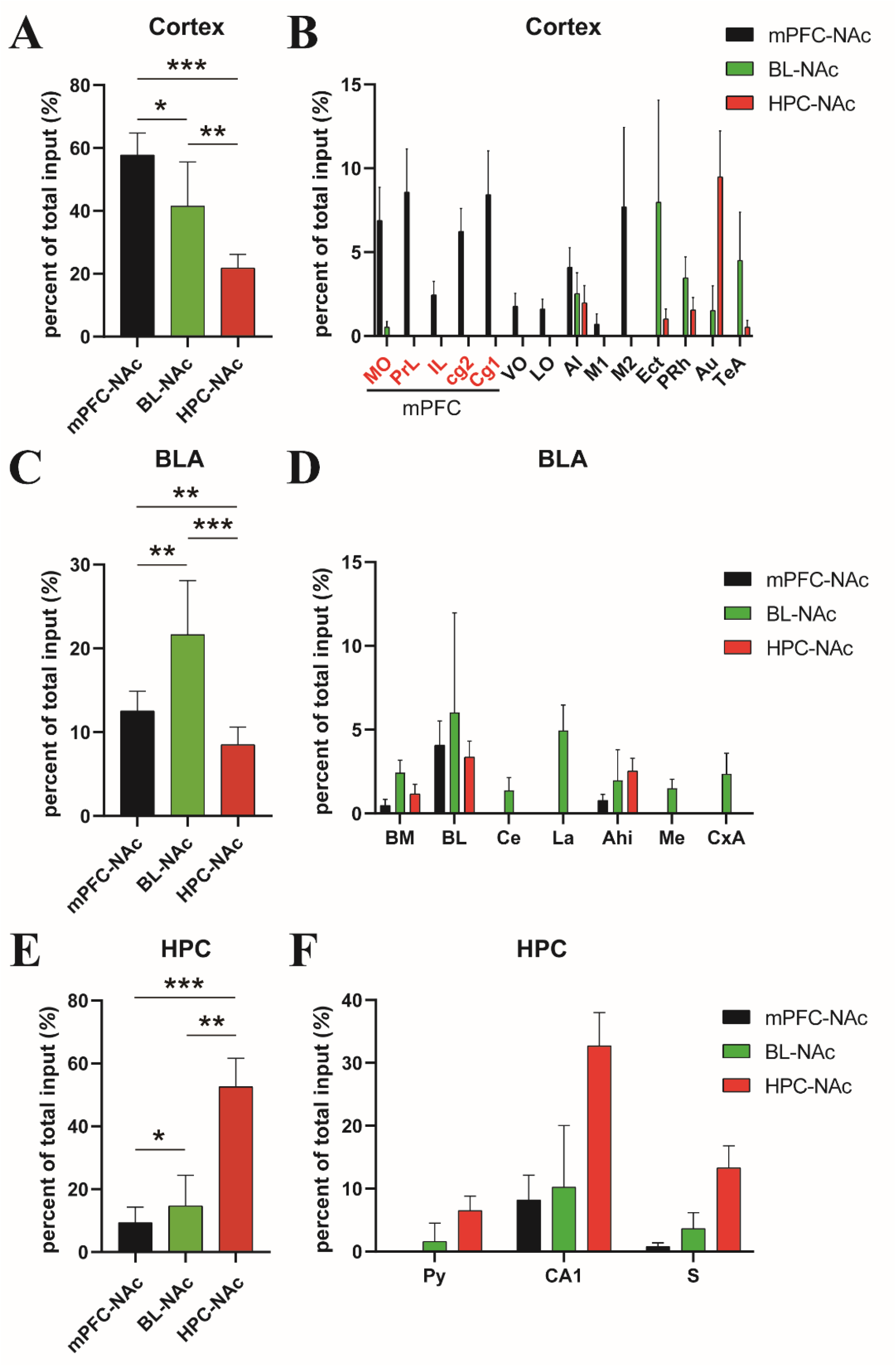
Quantitative analysis local input of mPFC/BL/HPC-NAc projectors. A. Comparative distribution of cortex input to mPFC/BL/HPC-NAc projectors. B. Input pattern of cortex to mPFC/BL/HPC-NAc projectors. C. Comparative distribution of amygdala input to mPFC/BL/HPC-NAc projectors. D. Input pattern of amygdala to mPFC/BL/HPC-NAc projectors. E. Comparative distribution of hippocampus input to mPFC/BL/HPC-NAc projectors. F. Input pattern of hippocampus to mPFC/BL/HPC-NAc projectors.

Amygdala provided more afferent inputs to BL-NAc projectors than to mPFC-NAc and HPC-NAc projectors (**Figure 7C**). BL-NAc projectors received disperse inputs from multiple subregions in the amygdala, including BL, Basomedial amygdaloid nucleus (BM), Central amygdaloid nucleus (Ce), Lateral amygdaloid nucleus (La), amygdalohippocampal area (AHi), medial amygdaloid nucleus (Me) and cortex-amygdala transition zone (CxA), while only AHi, BL and BM provided afferent inputs to HPC-NAc and mPFC-NAc projectors (**Figure 7D**).

HPC provided more afferent inputs to HPC-NAc projectors than to mPFC-NAc and BL-NAc projectors (**Figure 7E**). CA1 provided the most afferent inputs, and both of BL-NAc and HPC-NAc projectors received projections from Py, CA1 and S, thereinto, (**Figure 7F**).

It is note that mPFC/BL/HPC-NAc projectors received a high level of local inputs. Specifically, 32.62%±4.38% of inputs to mPFC -NAc projector were provided by mPFC, 6.04%±0.67% of inputs to BL-NAc projector were provided by BL, 32.72%±4.48% of inputs to HPC-NAc projector were provided by HPC (**Figure 7B,D,F**).

## Discussion

Previous study has revealed organization of input and output of NAc core and shell^11,30^, and some monosynaptic inputs of NAc projectors were traced by using TRIO method as well^31^. It has been found that NAc received a high input weight from mPFC, BL and HPC^4^. The circuits formed by these three brain regions and NAc participate in extensive and important brain functions. However, systematical analysis of the top-down afference of NAc in the range of the whole brain remains unclear. In this study, we used TRIO strategy to trace, manually count more than 550 thousand neurons throughout 114 brain regions and quantitatively analyze the whole brain input distribution of mPFC/BL/HPC-NAc projectors.

In our study, we found some arrestive top-down afferences of NAc. For example, Ent contributes inputs with a high weight to HPC-NAc projectors. It is noted that all of Ent, HPC and NAc contains neurons controlling mouse spatial navigation^4,32,33^.

Neurons in the Ent provide a map-like representation of the external spatial world^34^; hippocampus plays important roles in three strategies using for spatial navigation: path integration, stimulus-response association and map-based navigation^33^; lesion of NAc shell specifically impairs the acquisition of conditioned place preference and the use of spatial information^11^. Based on these results, we propose that Ent, HPC and NAc shell constitute a key circuit of spatial navigation.

Previous studies have shown that NAc received projections from HPC with a high input weight^4^. Our study found that more than half of afferent inputs of HPC-NAc projectors were from HPC, which suggesting that HPC not only directly regulate the activity of NAc, but also form local synaptic connections to indirectly regulate the activity of NAc through HPC-NAc projectors. The significance and operating mechanism of this dual regulation need to be further studied.

It is reported that most of the afferent inputs to mPFC were concentrated in the cortex. In our study, we found that it is cortex that provides most input to mPFC-NAc projectors. BLA receives local projections with a low weight^35^, while our results showed BL-NAc projectors receives projections from BLA with a high weight. Striatum provides important inputs to mPFC and BLA^12,35^, while our results found that few afferent inputs to mPFC/BL-NAc projectors come from striatum. Additionally, we have reported that the striatum provides few inputs to NAc^4^. If and how the striatum regulates NAc remains indistinct.

Previous study has reported that the MD is widely associated with cognition and receives information from association cortex^12^. In our study, we found that of the 14 subregions throughout the thalamus projecting to mPFC-NAc projectors, MD contribute the most of the inputs. Whether the circuits formed by thalamus and mPFC-NAc projectors is related to cognition needs further study.

## DATA AVAILABILITY

All data generated or analyzed in this study are included in the manuscript.

## ETHICS STATEMENT

The animal study was reviewed and approved by the Institutional Animal Care and Use Committee of Nanchang University

## AUTHOR CONTRIBUTIONS

Tonghui Xu, Jialiang Wu, Zhilong Chen and Zhao Li conceived and designed the project. Jialiang Wu completed all the virus injections experiment. Jialiang Wu, Zhilong Chen, Xueqi Gong performed data analysis. Jialaing Wu, Zhilong Chen and Xueqi Gong generated all Figures. Jialiang Wu, Tonghui Xu and Xueqi Gong wrote the paper.

## ACKNOWLEDGMENTS

We thank the Institute of Life Science, Nanchang University, Department of Laboratory Animal Science, Fudan University for their support with the equipment.

## REFERENCES

1 Groenewegen, H. J., Wright, C. I., Beijer, A. V. & Voorn, P. Convergence and segregation of ventral striatal inputs and outputs. Ann N Y Acad Sci 877, 49–63, doi:10.1111/j.1749-6632.1999.tb09260.x (1999).

2 Carlezon, W. A., Jr. & Thomas, M. J. Biological substrates of reward and aversion: a nucleus accumbens activity hypothesis. Neuropharmacology 56 Suppl 1, 122–132, doi:10.1016/j.neuropharm.2008.06.075 (2009).

3 Ito, R., Robbins, T. W., Pennartz, C. M. & Everitt, B. J. Functional interaction between the hippocampus and nucleus accumbens shell is necessary for the acquisition of appetitive spatial context conditioning. J Neurosci 28, 6950–6959, doi:10.1523/jneurosci.1615-08.2008 (2008).

4 Li, Z. et al. Cell-Type-Specific Afferent Innervation of the Nucleus Accumbens Core and Shell. Front Neuroanat 12, 84, doi:10.3389/fnana.2018.00084 (2018).

5 O’Connor, E. C. et al. Accumbal D1R Neurons Projecting to Lateral Hypothalamus Authorize Feeding. Neuron 88, 553–564, doi:10.1016/j.neuron.2015.09.038 (2015).

6 Nie, X. et al. Subregional Structural Alterations in Hippocampus and Nucleus Accumbens Correlate with the Clinical Impairment in Patients with Alzheimer’s Disease Clinical Spectrum: Parallel Combining Volume and Vertex-Based Approach. Front Neurol 8, 399, doi:10.3389/fneur.2017.00399 (2017).

7 Lüscher, C. The Emergence of a Circuit Model for Addiction. Annu Rev Neurosci 39, 257–276, doi:10.1146/annurev-neuro-070815-013920 (2016).

8 Russo, S. J. et al. The addicted synapse: mechanisms of synaptic and structural plasticity in nucleus accumbens. Trends Neurosci 33, 267–276, doi:10.1016/j.tins.2010.02.002 (2010).

9 Francis, T. C. & Lobo, M. K. Emerging Role for Nucleus Accumbens Medium Spiny Neuron Subtypes in Depression. Biol Psychiatry 81, 645–653, doi:10.1016/j.biopsych.2016.09.007 (2017).

10 Zahm, D. S. & Brog, J. S. On the significance of subterritories in the “accumbens” part of the rat ventral striatum. Neuroscience 50, 751–767, doi:10.1016/0306-4522(92)90202-d (1992).

11 Floresco, S. B. The nucleus accumbens: an interface between cognition, emotion, and action. Annu Rev Psychol 66, 25–52, doi:10.1146/annurev-psych-010213-115159 (2015).

12 Ährlund-Richter, S. et al. A whole-brain atlas of monosynaptic input targeting four different cell types in the medial prefrontal cortex of the mouse. Nat Neurosci 22, 657–668, doi:10.1038/s41593-019-0354-y (2019).

13 Pinto, L. & Dan, Y. Cell-Type-Specific Activity in Prefrontal Cortex during Goal-Directed Behavior. Neuron 87, 437–450, doi:10.1016/j.neuron.2015.06.021 (2015).

14 Bechara, A., Tranel, D. & Damasio, H. Characterization of the decision-making deficit of patients with ventromedial prefrontal cortex lesions. Brain 123 (Pt 11), 2189–2202, doi:10.1093/brain/123.11.2189 (2000).

15 Euston, D. R., Gruber, A. J. & McNaughton, B. L. The role of medial prefrontal cortex in memory and decision making. Neuron 76, 1057–1070, doi:10.1016/j.neuron.2012.12.002 (2012).

16 Laubach, M., Caetano, M. S. & Narayanan, N. S. Mistakes were made: neural mechanisms for the adaptive control of action initiation by the medial prefrontal cortex. J Physiol Paris 109, 104–117, doi:10.1016/j.jphysparis.2014.12.001 (2015).

17 Ridderinkhof, K. R., van den Wildenberg, W. P., Segalowitz, S. J. & Carter, C. S. Neurocognitive mechanisms of cognitive control: the role of prefrontal cortex in action selection, response inhibition, performance monitoring, and reward-based learning. Brain Cogn 56, 129–140, doi:10.1016/j.bandc.2004.09.016 (2004).

18 Schiff, H. C. et al. An Insula-Central Amygdala Circuit for Guiding Tastant-Reinforced Choice Behavior. J Neurosci 38, 1418–1429, doi:10.1523/jneurosci.1773-17.2017 (2018).

19 Yu, K., Garcia da Silva, P., Albeanu, D. F. & Li, B. Central Amygdala Somatostatin Neurons Gate Passive and Active Defensive Behaviors. J Neurosci 36, 6488–6496, doi:10.1523/jneurosci.4419-15.2016 (2016).

20 Yu, K. et al. The central amygdala controls learning in the lateral amygdala. Nat Neurosci 20, 1680–1685, doi:10.1038/s41593-017-0009-9 (2017).

21 Douglass, A. M. et al. Central amygdala circuits modulate food consumption through a positive-valence mechanism. Nat Neurosci 20, 1384–1394, doi:10.1038/nn.4623 (2017).

22 Kesner, R. P. & Rolls, E. T. A computational theory of hippocampal function, and tests of the theory: new developments. Neurosci Biobehav Rev 48, 92–147, doi:10.1016/j.neubiorev.2014.11.009 (2015).

23 Rolls, E. T. The storage and recall of memories in the hippocampo-cortical system. Cell Tissue Res 373, 577–604, doi:10.1007/s00441-017-2744-3 (2018).

24 Christakou, A., Robbins, T. W. & Everitt, B. J. Prefrontal cortical-ventral striatal interactions involved in affective modulation of attentional performance: implications for corticostriatal circuit function. J Neurosci 24, 773–780, doi:10.1523/jneurosci.0949-03.2004 (2004).

25 Piantadosi, P. T., Yeates, D. C. M., Wilkins, M. & Floresco, S. B. Contributions of basolateral amygdala and nucleus accumbens subregions to mediating motivational conflict during punished reward-seeking. Neurobiol Learn Mem 140, 92–105, doi:10.1016/j.nlm.2017.02.017 (2017).

26 Britt, J. P. et al. Synaptic and behavioral profile of multiple glutamatergic inputs to the nucleus accumbens. Neuron 76, 790–803, doi:10.1016/j.neuron.2012.09.040 (2012).

27 Sosa, M., Joo, H. R. & Frank, L. M. Dorsal and Ventral Hippocampal Sharp-Wave Ripples Activate Distinct Nucleus Accumbens Networks. Neuron 105, 725–741.e728, doi:10.1016/j.neuron.2019.11.022 (2020).

28 Schwarz, L. A. et al. Viral-genetic tracing of the input-output organization of a central noradrenaline circuit. Nature 524, 88–92, doi:10.1038/nature14600 (2015).

29 Osakada, F. et al. New rabies virus variants for monitoring and manipulating activity and gene expression in defined neural circuits. Neuron 71, 617–631, doi:10.1016/j.neuron.2011.07.005 (2011).

30 Ma, L., Chen, W., Yu, D. & Han, Y. Brain-Wide Mapping of Afferent Inputs to Accumbens Nucleus Core Subdomains and Accumbens Nucleus Subnuclei. Front Syst Neurosci 14, 15, doi:10.3389/fnsys.2020.00015 (2020).

31 Loureiro, M. et al. Social transmission of food safety depends on synaptic plasticity in the prefrontal cortex. Science 364, 991–995, doi:10.1126/science.aaw5842 (2019).

32 Kunz, L. et al. Mesoscopic Neural Representations in Spatial Navigation. Trends Cogn Sci 23, 615–630, doi:10.1016/j.tics.2019.04.011 (2019).

33 Jin, W., Qin, H., Zhang, K. & Chen, X. Spatial Navigation. Adv Exp Med Biol 1284, 63–90, doi:10.1007/978-981-15-7086-5_7 (2020).

34 Butler, W. N., Hardcastle, K. & Giocomo, L. M. Remembered reward locations restructure entorhinal spatial maps. Science 363, 1447–1452, doi:10.1126/science.aav5297 (2019).

35 Fu, J. Y. et al. Whole-Brain Map of Long-Range Monosynaptic Inputs to Different Cell Types in the Amygdala of the Mouse. Neurosci Bull 36, 1381–1394, doi:10.1007/s12264-020-00545-z (2020).

