## Supplementary Table 1, Supplementary Figure 1 for "Whole-Brain Map of Top-down afferent Inputs to Nucleus Accumbens of the Mouse"

Tonghui Xu^1*^，Jialiang Wu^2#^, Zhilong Chen^3,4#^, Zhao Li^5#^

^1^Department of Laboratory Animal Science, Fudan University, Shanghai 200000, China.

^2^Institute of Life Science, Nanchang University, Nanchang 330031, China.

^3^Key Laboratory of Biomaterials of Guangdong Higher Education Institutes, Department of Biomedical Engineering, Jinan University, Guangzhou 510632, China

^4^Piedmont Medical Technology Co., Ltd., Zhuhai 519031, China

^5^Reproductive Medicine Center, Tongji Hospital, Tongji Medical College, Huazhong University of Science and Technology, Wuhan, People's Republic of China.

^#^These authors contributed equally to this work

**ABSTRACT**

Nucleus accumbens (NAc) is an important part of basal ganglia and receives major transsynaptic inputs from medial prefrontal cortex (mPFC), basolateral amygdala nucleus (BL) and hippocampus (HPC). The neural circuits formed by mPFC, BL, HPC with NAc are closely related to diverse brain functions such as decision-making, social behavior, reward seeking behavior and aggressive behavior. However, a lack of information about network structure of NAc top-down afference has limited our understanding of mechanisms of these functions. Here, we systematically analyzed whole brain patterns of inputs to mPFC/HPC/BL-NAc projectors via using (TRIO) labeling strategy. We found that the upstream input patterns between mPFC-core and mPFC-shell projections, BL-core and BL-shell projections, HPC-core and HPC-shell projections were similar. The projections from mPFC and BL to NAc receive a wide range of inputs, while the projections from HPC to NAc receive convergent inputs. Furthermore, all of mPFC/HPC/BL-NAc projectors receive considerable inputs from respective local regions. This study lays a foundation for further analysis of complex NAc functional mechanisms.

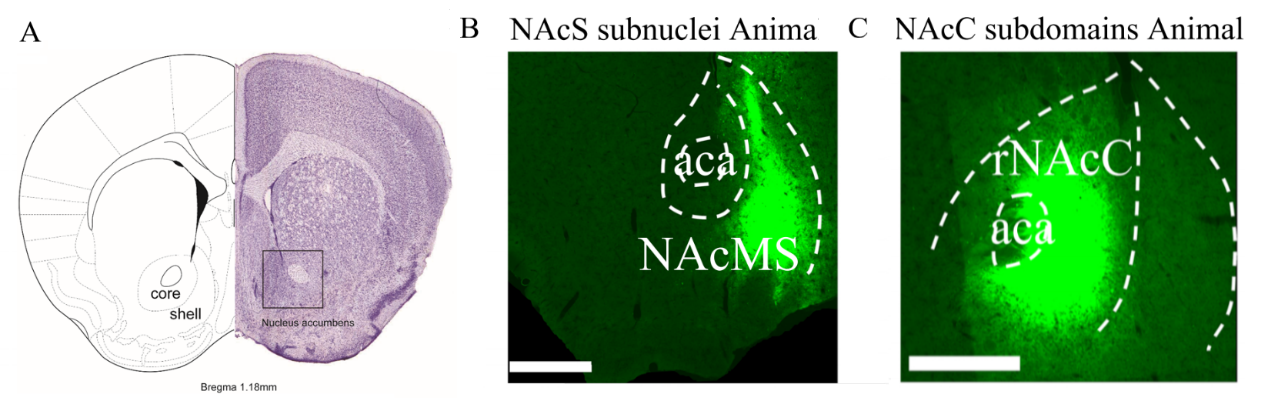

**Fig. S1 | NAc core and shell injection sites.**

A. Anatomical location and sub-regional division of the NAc in mouse brain.

B. Representative images of NAc shell injection sites.

C. Representative images of NAc core injection sites.

**Supplementary Table 1 Abbreviation for brain area**

| **Abbreviation for brain area** | |
| --- | --- |
| Agranular insular area/Gustatory | AI/GU |
| Anterior cingulate area | ACA |
| Anteromedial nucleus of thalamus | AM |
| Anteromedial visual area | VISl |
| Auditory areas | AU |
| Basolateral amygdala nucleus | BL |
| Basomedial amygdaloid nucleus | BM |
| Central amygdaloid nucleus | Ce |
| Cingulate cortex | Cg |
| Central lateral nucleus of thalamus | CL |
| Central medial nucleus of thalamus | CM |
| Cerebellar nuclei | CBN |
| Claustrum | CLA |
| Contralateral dentate nucleus/Interposed nucleus | con-DN/IP |
| Contralateral primary motor area | con-MOp |
| Contralateral secondary motor area | con-MOs |
| Cortical plate | CTXpl |
| Cortical subplate | CTXsp |
| Diagonal band nucleus | NDB |
| Dorsal nucleus raphe | DR |
| Dorsal peduncular area/Taenia tecta | DP/TT |
| Extranasal cortex | Ect |
| Entorhinal area | ENT |
| Gigantocellular/Intermediate reticular nucleus | GRN/IRN |
| Hippocampus | HPC |
| Infralimbic area | IL |
| Lateral amygdaloid nucleus | La |
| Lateral posterior nucleus of thalamus | LP |
| Lateral visual area | VISl |
| Primary motor cortex | M1 |
| Secondary motor cortex | M2 |
| Mediodorsal nucleus of thalamus | MD |
| Medial septal nucleus | MS |
| Medulla | MY |
| Midbrain, behavioral state related | MBsta |
| Midbrain, motor related | MBmot |
| Medial orbital cortex | MO |
| Medial prefrontal cortex | mPFC |
| Midbrain reticular nucleus | MRN |
| Nucleus accumbens | NAc |
| Orbital area, lateral part | ORBl |
| Orbital area, ventrolateral part | ORBvl |
| Orbital area, medial part | ORBm |
| Pallidum | PAL |
| Pallidum, dorsal region | PALd |
| Paracentral nucleus of thalamus | PCN |
| Parafascicular nucleus of thalamus | PF |
| Piriform cortex | Pir |
| Pedunculopontine nucleus | PPN |
| Pons | P |
| Pontine gray | PG |
| Posterior complex of thalamus | PO |
| Perinasal cortex | PRh |
| Prelimbic area | PrL |
| Primary motor cortex | MOp |
| Primary somatosensory area, barrel field | SSp-bfd |
| Primary somatosensory area, lower limb/trunk | SSp-ll/tr |
| Primary somatosensory area, upper limb/mouth | SSp-ul/m |
| Temporal association cortex | TeA |
| Primary visual area | VISp |
| Retrosplenial area | RSP |
| Secondary motor cortex | MOs |
| Substantia innominata | SI |
| Substantia nigra, reticular part | SNr |
| Superior central nucleus raphe | CS |
| Superior colliculus | SC |
| Supplemental somatosensory area | SSs |
| Thalamus | TH |
| Ventral anterior-lateral complex of thalamus | VAL |
| Ventral medial nucleus of thalamus | VM |
| Ventral orbital cortex | VO |
| Ventral tegmental area | VTA |
